# Application of Large Language Models for Annotating Genes into Reactome Pathways

**DOI:** 10.64898/2025.12.20.695723

**Authors:** Guanming Wu, Lisa Matthews, Nathan Boyer, Marija Milacic, Deidre Beavers, Nancy T Li, Bruce May, Karen Rothfels, Veronica Shamovsky, Ralf Stephan, Marc Gillespie, Henning Hermjakob, Peter D’Eustachio, Lincoln Stein

## Abstract

Reactome is the most comprehensive, open source, open access biological pathway knowledgebase, widely used in the research community. To ensure the highest quality of its content, human pathway data in Reactome is manually curated. However, manual curation is labor-intensive, time-consuming, and increasingly difficult to keep up with the ever-growing biomedical literature. Large language model (LLM)–driven artificial intelligence (AI) technologies are transforming many fields, including bioinformatics resource development. Applying LLM/AI technologies in Reactome may offer a powerful way to scale curation and consolidate pathway-related data into a single resource. This manuscript describes the first stage of our attempt to adopt LLM/AI technologies for Reactome manual curation. We developed an LLM workflow that can assist curators in adding new genes to existing pathways and refining the functional annotations of existing ones. The workflow predicts pathways in which genes are likely to function, identifies PubMed-indexed literature that may support these predictions, generates text summaries describing potential molecular mechanisms, and extracts functional relationships among biological entities from full-text PDF papers. To validate the workflow output, we used a computational approach based on semantic similarity between LLM workflow-generated summaries and Reactome manual annotations. The results show significant enrichment of high-similarity matches. Manual evaluation of 19 genes indicated that more than half of the outputs are useful for supporting curation. Based on these results, we developed an enhanced workflow that incorporates protein-protein interaction data, facilitating Reactome’s reaction-based annotation. In summary, our initial adoption of LLM/AI technologies produced encouraging results and provides a practical framework for integrating AI-assisted methods into Reactome’s curation pipeline. The strategies described here may be broadly applicable to community knowledgebases in general.

## Introduction

Biological pathways are foundational to modern biological research. They provide structured frameworks for organizing discoveries, elucidating molecular and cellular mechanisms underlying phenotypes, and guiding the development of new therapies for cancer and other complex diseases. Pathways also underpin systems biology and are indispensable for large-scale omics analyses, enabling researchers to integrate scattered molecule-level observations, uncover hidden relationships among genes, proteins and other types of molecules, and generate experimentally testable hypotheses. Biological pathway databases are resources where pathways are organized systematically and comprehensively, serving as essential knowledgebases supporting a wide range of research activities^1^.

Reactome is one of the most widely used open-source, open-access biological pathway knowledgebases, providing curated pathway data, together with analysis tools and visualization capabilities, for the biomedical research community to perform pathway-based data analysis and visualization^2^. To ensure the highest data quality, content in Reactome is manually curated from publicly available, PubMed-indexed literature. Although this approach has produced a highly trusted resource, the manual curation process is labor-intensive, time-consuming, and increasingly difficult to scale in the face of rapid growth in biological data and publications that are driven by high-throughput technologies at the individual cell level.

Large language models (LLMs)-driven artificial intelligence (AI) technologies are transforming many fields, including biomedical research^3^. LLMs are trained on extensive bodies of human-written text, demonstrating strong capabilities in understanding and synthesizing complex knowledge. High-quality pathway databases like Reactome are fundamentally literature-driven, and represent a natural and promising domain for the application of LLM-based AI technologies. However, early adoption revealed substantial challenges, most notably hallucinations^4, 5^: LLMs can generate nonexistent references, unsupported assertions, and incorrect mechanistic statements. Retrieval-augmented generation (RAG)^6^ mitigates this issue by grounding LLM outputs in user-provided contextual information, substantially reducing hallucination, though not eliminating it entirely.

To address the bottleneck of manual curation while maintaining high data quality, we are integrating LLM-based AI technologies into Reactome’s curation workflow to develop a curator-in-the-loop LLM application. This manuscript reports the first step in this effort: developing an LLM workflow to assist curators in annotating human genes into existing Reactome pathways. The workflow predicts pathways potentially relevant to a query gene, identifies supporting literature from PubMed by analyzing abstracts using semantic similarity, and generates summary text using LLMs to facilitate manual review by Reactome curators. Our workflow adopts RAG extensively to reduce the chance of hallucination. Insights from this initial stage guide refinement of the approach and support the development of a practical AI-assisted system capable of scaling Reactome’s manual curation without compromising accuracy.

## Results

### Overall workflow of using LLMs for annotating genes into Reactome pathways

To reduce the risk of hallucinations by LLMs, we incorporated RAG extensively throughout the workflow (**Figure 1**). The workflow consists of several steps aimed at annotating the function of a query gene, whether or not it has been annotated in Reactome, into existing Reactome pathways based on information derived from both external resources and internal Reactome data. First, the workflow queries PubMed for papers related to the query gene, with an emphasis on its interactions, reactions, or pathways. For the example gene TANC1, the query uses the keywords “TANC1 interactions,” “TANC1 reactions,” or “TANC1 pathways,” and retrieves the abstracts of the top-ranked papers. The workflow also queries the Reactome IDG portal through its RESTful API deployed at idg.reactome.org^7, 8^. The portal was developed to place understudied human proteins in the context of Reactome pathways and to support the generation of experimentally testable hypotheses. It uses functional interactions (FIs)^9^ between the query gene and Reactome-annotated genes to infer interacting pathways for the query genes. The FIs were predicted using a random forest model trained on 106 protein/gene pairwise relationship features that collectively cover the whole human proteome^8^. These interacting pathways are candidate pathways into which the query gene may be annotated.

**Figure 1.**
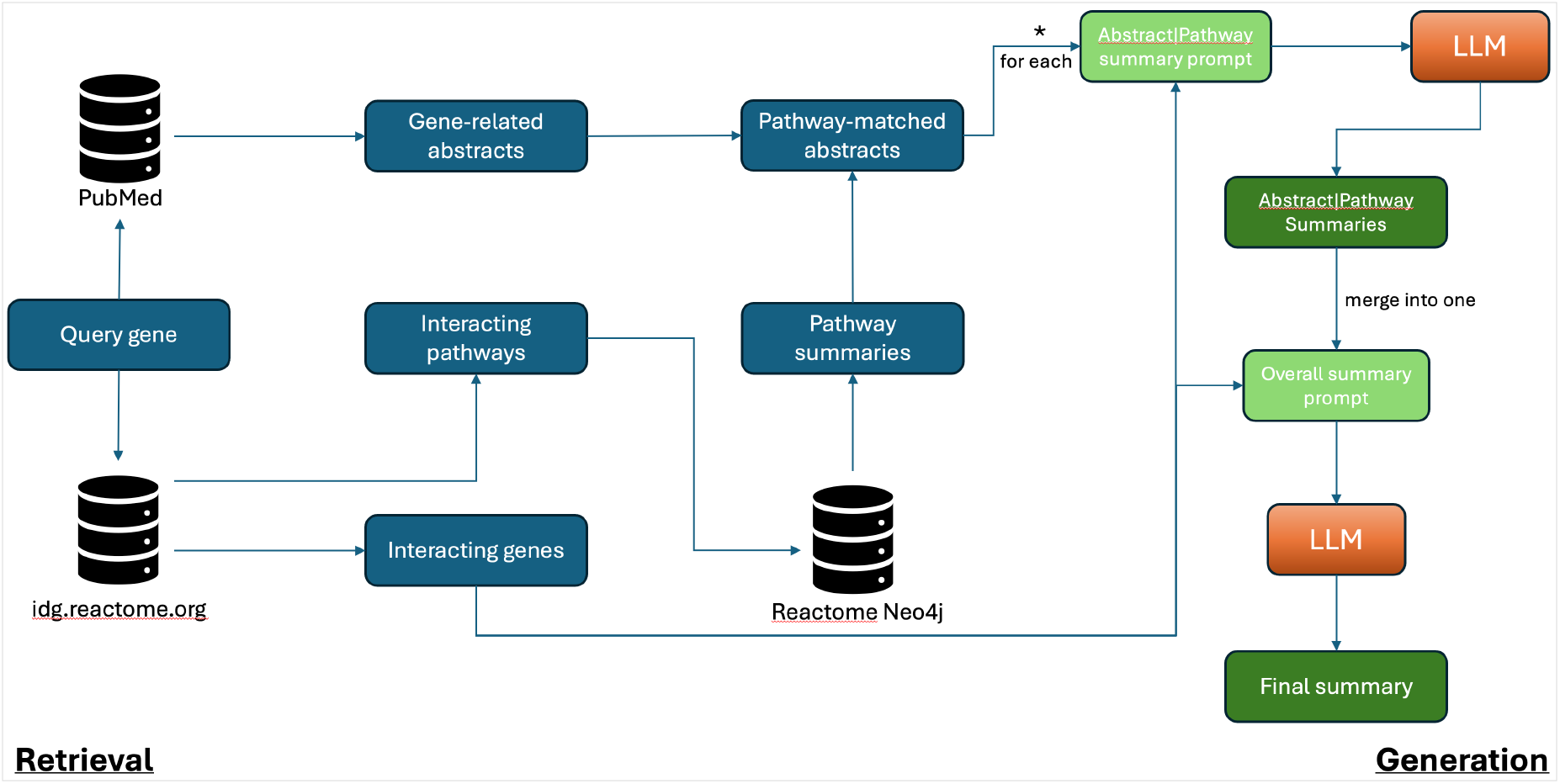
The LLM workflow for annotating human genes into Reactome pathways.

The retrieved abstracts are compared with the text descriptions of interacting pathways by first embedding both the abstracts and the summary texts of pathways fetched from the Reactome Neo4j database^10^ into numeric vectors, and then calculating semantic similarity. The resulting similarity scores reflect the likelihood that the papers are relevant to the interacting pathways, which may support the annotation of the query gene in the interacting pathways. Matched pathways and abstracts are then summarized into text using the OpenAI API. To generate the final output for curators, the list of all summaries for abstract–pathway text pairs is further condensed using the API. To ground the summaries and further reduce the chance of hallucination, genes that functionally interact with the query gene and are annotated in the interacting pathways are supplied to the API as a part of contextual information. This additional grounding helps guide the LLM’s attention to mentions of these interacting genes and facilitates downstream molecular mechanism–based annotation in Reactome.

### Validation of the LLM generated summaries via a computational approach

To validate the output of the LLM workflow for Reactome pathway curation, we developed a computational approach that compares annotations predicted by the LLM workflow with existing curators’ manual annotations for a set of genes that had not been annotated in an earlier Reactome release (the baseline release) but were annotated in a later release and therefore serve as validation data. The LLM workflow generates text annotations for these genes based on the content present in the database at the baseline release, and the results are compared against manually curated annotations for those same genes in the later release. To enable semantic comparison, we use LLMs to generate summaries of the curated content for each query gene, then embed both the LLM-predicted summaries and the curator-annotated summaries into numeric vectors for cosine similarity analysis (**Figure 2A**).

**Figure 2.**
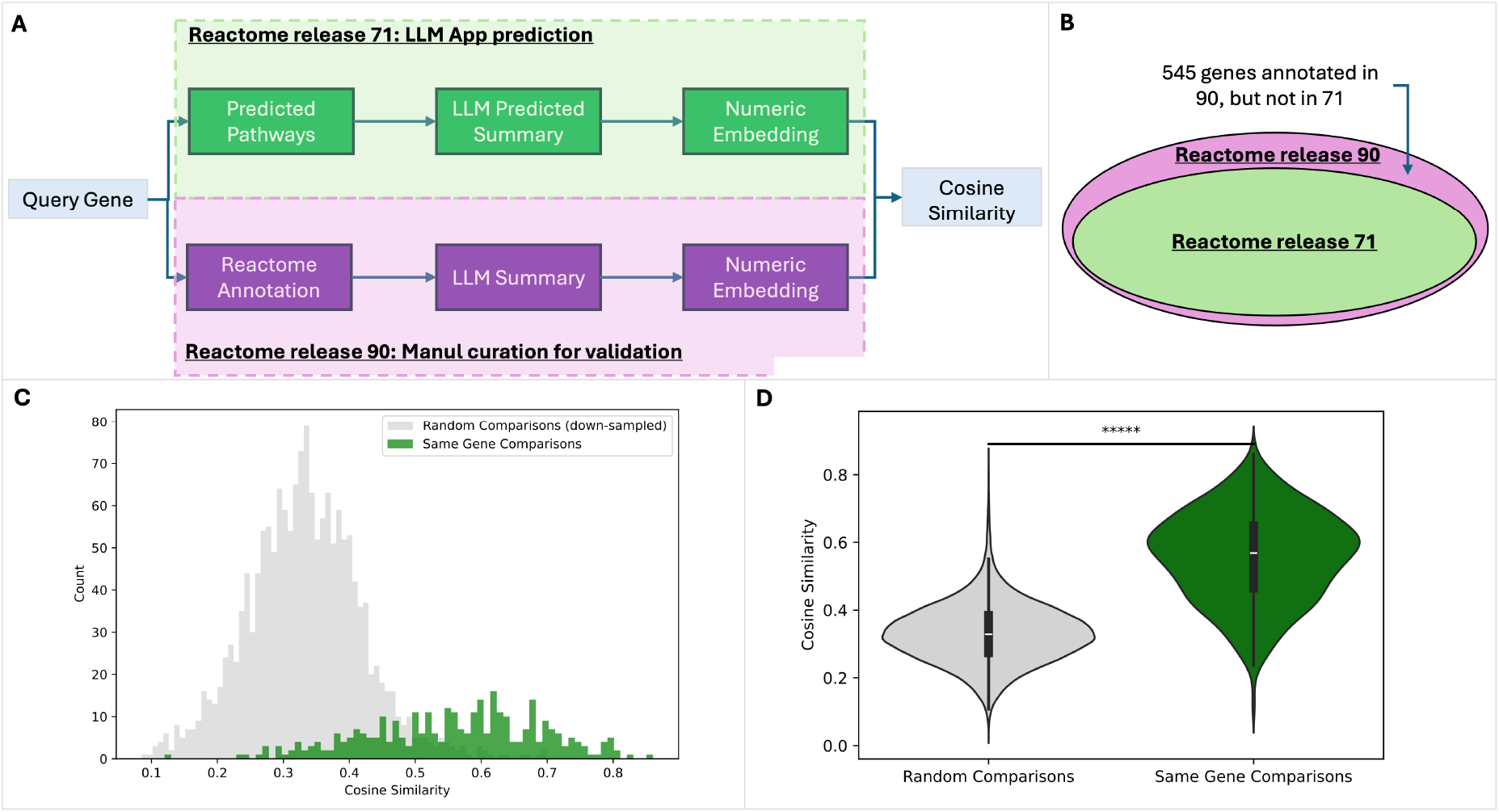
Computational approach for validating the LLM workflow generated pathway annotations for query genes by using genes that were annotated in Reactome release 90 but not in release 71. Semantic similarity was calculated between LLM workflow generated annotations and curators’ manual annotations. **A**. Workflow of the computational validation approach. **B**. Number of genes annotated in release 90 but not in release 71. (Note: Ellipse areas are not proportional to gene counts.). **C**. Histogram showing the right-tailed distribution of semantic similarities between LLM-generated pathway annotations and curators’ manual annotations (green), compared against a random background (gray). **D**. Violin plot comparing the distributions of semantic similarity scores for the random background and the LLM-generated annotations (p-value = 1.49E-150 based on Mann-Whitney U Test).

**Figure 3.**
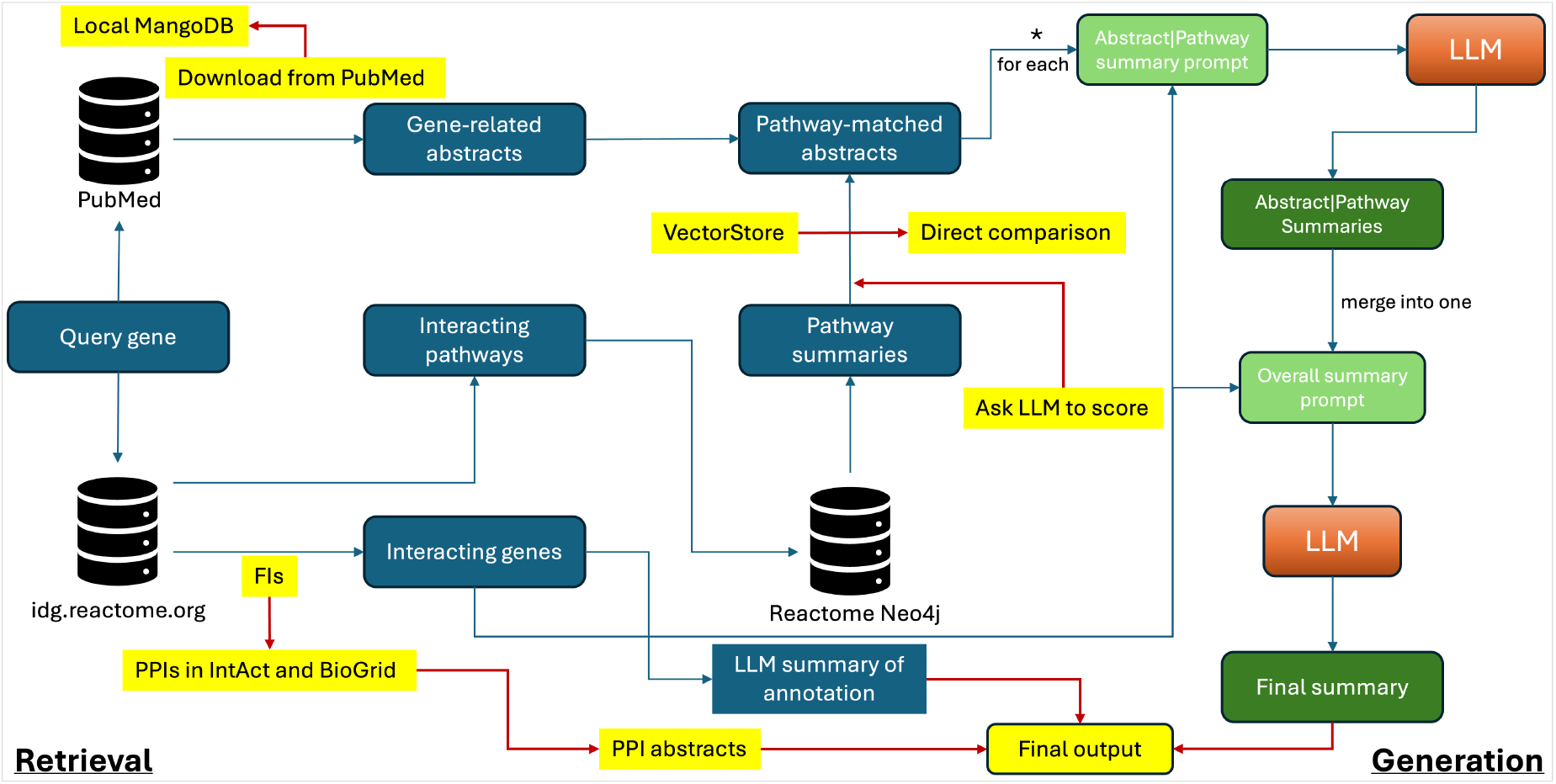
The enhanced LLM workflow to address the drawbacks discovered from the manual validation. Features in yellow are related to enhancements.

We selected Reactome release 71 as the baseline, which was also used as the positive training dataset for the random forest model that predicts functional interactions^8^, the basis for inferring interacting pathways for query genes, and chose release 90 as the validation set. In total, we identified 545 human genes that were not annotated in release 71 but annotated in release 90 (**Figure 2B**). Of these, the LLM workflow generated text summaries for 326 genes (∼60%). Some of the remaining 545 genes lacked functional interaction partners in annotated pathways (e.g., genes whose protein products act solely as catalysts in small-molecule reactions). Consequently, the validation workflow failed to generate text summaries for these genes, due to its focus on functional relationships among proteins.

Semantic similarity analysis of the 326 LLM-summarized genes showed a right-shifted distribution, indicating significantly higher similarity compared with the random background generated by permuted comparisons between LLM-predicted text and LLM-summarized curators’ manual annotations for the same 326 genes (**Figure 2C** and **2D**, p-value = 1.49E-150 based on Mann-Whitney U Test). This result implies that the pathway annotations generated by the LLM workflow for the query genes are significantly semantically similar to Reactome curators’ manual annotations, providing a foundation for the LLM workflow as a tool to support Reactome curation.

### Manual validation of the LLM generated summaries for Reactome curation

To further validate the output of the LLM workflow, we hand-picked 19 genes that Reactome curators were actively working on in their individual curation projects and asked the curators to record the time they spent using the LLM workflow to obtain results, validate those results by checking PubMed or other resources such as UniProt^11^, IntAct^12^, and the Gene Ontology^13, 14^, and document their observations by manually evaluating the output. Overall, the LLM workflow produced good or acceptable results for 11 of the 19 genes (58%) with reasonable use of human curator time (less than 40 minutes), giving us confidence that the LLM workflow is moving in the right direction to provide an AI-powered, curator-in-the-loop tool to scale Reactome’s manual curation (**Table 1**. For a detailed version of this table, see https://github.com/reactome/curator-tool-llm/blob/main/validation/LLM%20feature%20testing-Jan.2025.xlsx).

**Table 1.** Manual validation results for 19 hand-picked genes Reactome curators were annotating for their individual projects.

| Query gene | Comment | Score | Time spent |
| --- | --- | --- | --- |
| CHD2 | Found two related papers with a reasonable summary | good | 2 min |
| ETS1 | Found a related paper with a good summary. | good | n/a |
| PAPOLG | Good summary with related papers | good | 41 min |
| TECPR1 | Good summary with related papers | good | 15 min |
| ATG16L2 | Results focused on FI partners. Summary and papers are relevant. | ok | 20 min |
| ERP29 | Found useful papers with a reasonable summary. May miss some useful papers. | ok | 41 min |
| HES2 | Reasonable summary with related papers | ok | 20 min |
| HES6 | Found related papers. May overstate the role of the query gene in the adipocyte differentiation based on one paper. | ok | 20 min |
| PER3 | Reasonable summary with related papers. But may miss some useful papers. | ok | 41 min |
| SERPINF1 (PEDF) | Found useful papers with a reasonable summary. May miss some useful papers. | ok | 41 min |
| TECPR2 | Reasonable summary with related papers | ok | 20 min |
| BIK (BIP1) | The fetched paper is not relevant. | bad | 41 min |
| CHD1 | Focused on the query gene's FI partner, CHD4. Maybe over-reasoned. | bad | 7 min |
| ERC1 | Fetched papers focused on FI partners. Not directly related to the query gene. | bad | n/a |
| GATC | Fetched papers focused on FI partners. Not directly related to the query gene. | bad | 15 min |
| H2AB2, H2AB3 | No result is found though manual PubMed search found several useful papers. | bad | 41 min |
| HABP2 | FSAP is a synonym of HABP2, the query gene. The LLM cannot see they are the same gene and treated FSAP as an interactor of HABP2. | bad | n/a |
| HRK | Fetched papers focused on FI partners. Not directly related to the query gene. | bad | 10 min |
| NBL1 (DAN) | No result is found though manual PubMed search found several useful papers. | bad | 41 min |

Based on the curators’ observations, we identified several drawbacks in the first iteration of the LLM workflow with respect to its utility in supporting manual curation:

#### 1. Certain Interactions are not supported by PMIDs (e.g., ATG16L2, CHD1, ERC1, and GATC)

The LLM app leverages the Reactome IDG portal to predict interacting pathways for the query genes. The portal uses predicted functional interactions, which may not have strong experimental evidence even though their confidence is high based on the performance of the random forest model used for prediction. Because Reactome curation focuses on molecular mechanisms supported by published experimental evidence, FI-based predictions may have limited value in assisting manual curation if they are not supported by experimental evidence.

#### 2. The output may be too high-level, resembling pathway summaries rather than focusing on specific molecular mechanisms, and may miss relevant papers (e.g., ERP29)

For performance and data-accessibility reasons, the LLM workflow currently analyzes abstracts only, which may cause it to miss papers containing detailed mechanistic information that is more relevant to Reactome curation.

#### 3. Some matches between pathways and supporting papers are too vague or incorrect (e.g., BIK, HES6, and CHD1)

Although the LLM workflow provides extensive context to ground LLM reasoning, hallucinations may still occur.

#### 4. No results were found for some genes (e.g., H2AB2 and NBL1)

Some pre-set thresholds were used to fetch predicted functional interactions and interacting pathways for the query genes. These thresholds may have been set too high, preventing useful results from being retrieved. It is also possible that no experimental evidence has been published to support the predicted interacting pathways.

#### 5. Synonyms are sometimes not understood (e.g., HABP2)

This may reflect a limitation of the LLMs used, which may not reliably distinguish or map gene synonyms.

### Enhancement of the LLM workflow to support Reactome manual curation

To overcome the drawbacks identified through manual validation, we made several enhancements, including importing annotatable protein–protein interactions to filter functional interactions, introducing two approaches for abstract scoring to increase the likelihood of their relevance to inferred interacting pathways for the query gene, and improving the performance of downloading abstracts from PubMed.

The functional interactions used to predict interacting pathways for query genes were derived from 106 human protein/gene pairwise relationship features using a machine learning approach, many of which are supported by gene co-expression in tissues and cancers and may not have been experimentally validated. In contrast, protein–protein interaction (PPI) databases typically host experimentally validated PPIs that are manually curated. Reactome’s annotation of complexes and reactions is usually based on physical interactions between biological molecules. To better support Reactome’s manual curation of biochemical reaction–based molecular mechanisms, we imported manually curated, experimentally validated PPIs from two widely used PPI databases, IntAct^12^ and BioGRID^15^. All imported PPIs are supported by PubMed-indexed papers, providing an additional source of context for the LLM workflow when generating summaries for annotating genes into pathways.

During manual validation, we found that some abstracts were too vague to be useful or were not matched appropriately to support the inferred interacting pathways. To increase the relevance of abstracts, we implemented an approach to score them based on semantic similarity by directly calculating cosine similarity between the embedded abstracts and pathway summaries, instead of relying on the built-in vector store search. Additionally, we asked the LLM to act as a judge (LLM-judge)^16^ to evaluate the relevance of each abstract to the pathway. The final match between an abstract and a pathway is determined using both scores. Our testing indicated that this two-way approach resulted in higher relevance between abstracts and interacting pathways and filtered out false positive matches.

The LLM workflow relies heavily on abstracts downloaded from PubMed to identify relevant papers that can support manual annotation. Due to the enforced rate limit of fewer than 10 requests per second through NCBI’s E-utilities API, downloading abstracts became a bottleneck in the workflow. To overcome this limitation, we downloaded the PubMed annual baselines from the NCBI web site and loaded all abstracts into a local MongoDB instance, resulting in a substantial improvement in abstract-processing performance.

### The LLM app in the web-based Reactome curation tool

To support Reactome curators’ manual curation, we have implemented the LLM workflow as a web-based application in the upcoming new Reactome curation tool and deployed it at http://curator.reactome.org/curatortool/llm_apps_view. Curators can enter a query gene in the input panel, and open the configuration panel to choose the protein interaction sources used to predict interacting pathways for the query gene (**Figure 4A**). Currently, the LLM app supports human protein–protein interactions collected from IntAct and BioGRID only. Curators may choose whether to filter PPIs using Reactome FIs. They may also use Reactome FIs alone to predict interacting pathways, which can identify potential pathways that may not yet have experimental evidence supporting molecular mechanism–based annotation, as shown in the manual validation results above. In the configuration panel, curators can adjust thresholds for selecting Reactome FIs used for filtering PPIs, as well as for specifying the number of top abstracts returned from the PubMed query to be analyzed by the LLM app. Curators may also set thresholds based on semantic similarity and LLM scores measuring the relevance between abstracts and Reactome pathways, allowing them to filter out less relevant abstracts. Thresholds for top pathways and FDR are used to determine the total number of pathways included in the analysis.

**Figure 4.**
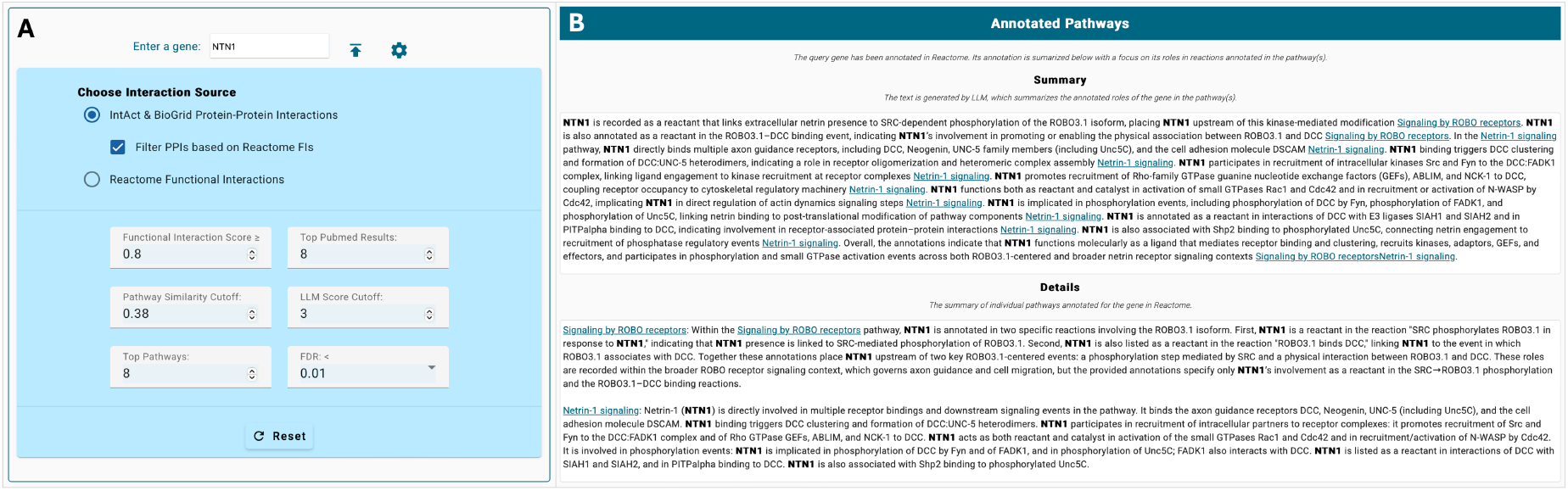
Screenshots of the LLM App in the web-based Reactome curation tool for query gene input, configuration and annotated pathways. **A**. The input panel where curators enter a query gene, along with the configuration panel used to choose the data sources for predicting interacting pathways and to set thresholds. **B**. The LLM-generated summary for manual annotation of a gene (e.g., NTN1) that has already been curated in Reactome.

The query gene (e.g., NTN1) may already be annotated in Reactome. For such genes, the LLM app generates an overall summary of Reactome’s existing annotation for the query gene. The app also provides detailed information about the specific biochemical reaction–based annotations in the details section. Each annotated pathway has its own subsection within this section, while the summary section provides an overview of all pathways (**Figure 4B**).

The main goal of the LLM app is to assist Reactome manual curation by identifying pathways into which the query gene may be annotated, along with published references that support the annotations. The predicted pathway section (**Figure 5A**) presents the predicted results for the query gene. The Summary subsection provides an overview of how the query gene may be functionally related to annotated Reactome pathways, supported by relevant literature. This overview focuses on molecular interactions between the query gene and its interaction partners in the target pathway. The Details subsection links PubMed literature (identified by PMIDs) to the target pathways. The LLM-generated text summaries provide succinct explanations of why and how each reference supports the annotation of the query gene in the corresponding pathway. For each PubMed reference– pathway pair, the app also displays the semantic similarity score and the LLM judge score, as well as the FDR value used to assess the significance of the predicted interacting pathway for the query gene. As shown in **Figure 5A**, a single pathway may be supported by multiple references.

**Figure 5.**
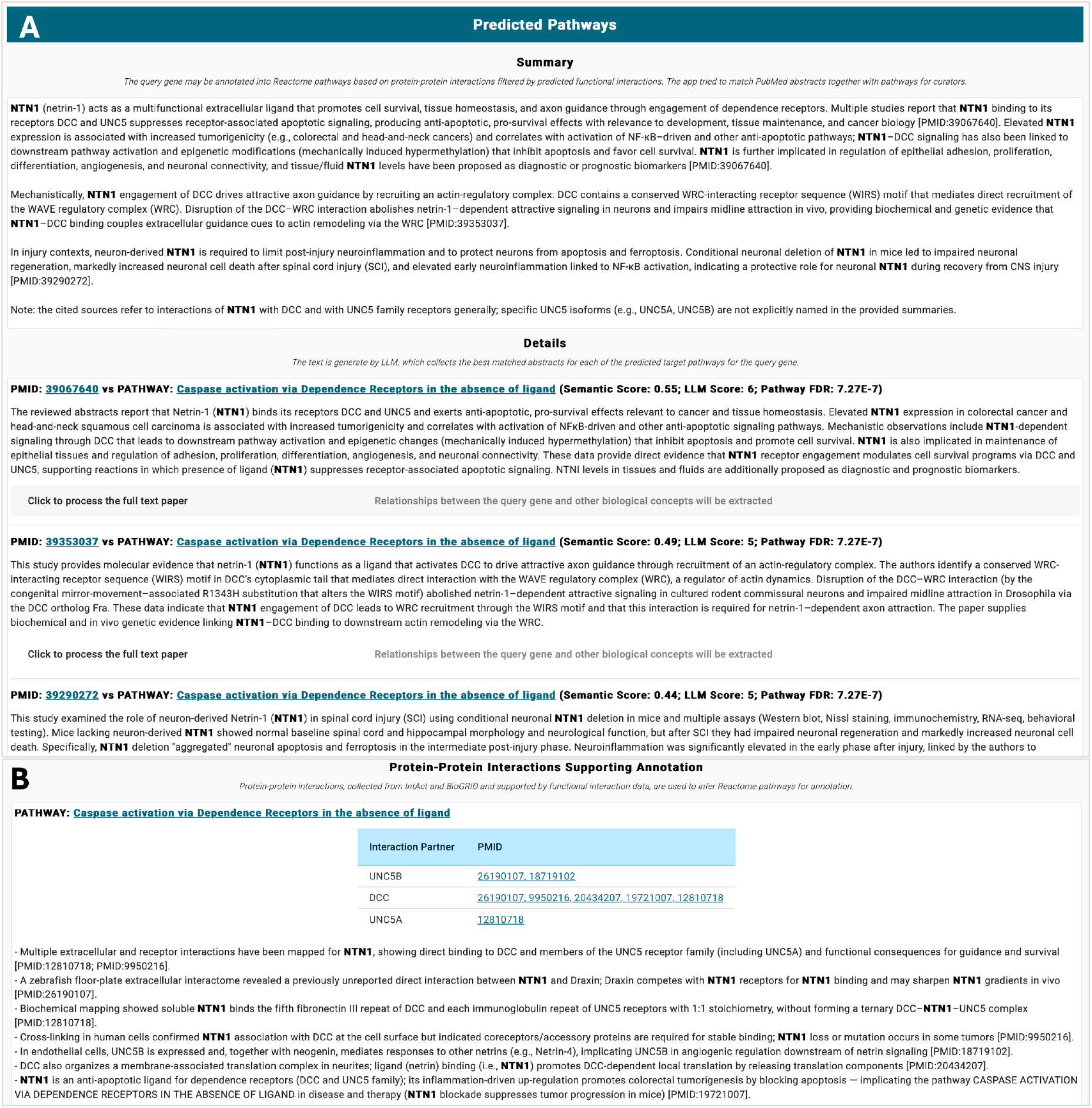
Screenshots of the LLM App in the web-based Reactome curation tool for predicted pathways and protein-protein interactions. **A**. Predicted pathways panel shows the summary and the details about the LLM workflow-predicted results for the query gene, including pathways it might be annotated into as well as the supporting literature PubMed identifiers (PMIDs). **B**. Summary for protein-protein interactions supporting the predicted results.

To provide annotatable references for Reactome manual curation, the original references used to annotate protein–protein interactions are also displayed for the target pathways (**Figure 5B**). The LLM app generates a summary of the interactions between the query gene and its interaction partners based on the abstracts of these original papers. This text may offer an additional type of evidence for annotating the query gene into the target pathway, in cases where these interaction references are not included in the Predicted Pathways subsection (**Figure 5A**).

Abstracts often contain limited descriptions of the molecular mechanisms required for Reactome manual curation. To address this, the LLM app includes a feature that allows curators to upload full-text PDF papers and extract functional relationships among biological entities (**Figure 6**). The extracted relationships, along with their original text sources, are listed under the section corresponding to the match between a PubMed reference and the target pathway. These relationships provide curation materials that align more closely with the Reactome data model and are better suited for potential future automated import into the Reactome database.

## Discussion

Knowledgebases like Reactome and other pathway databases, model organism databases, data resources like UniProt and ChEBI^17^, and ontologies like GO, must be comprehensive and accurate to be usable. These resources maintain teams of expert human curators to review published experimental data, extract, digest and then synthesize useful bits to generate reliable annotations. Even if funding for these resources were not limited, expert manual curation cannot readily be scaled to handle increasing numbers of publications and the massive amounts of data generated by high-throughput strategies like single-cell omics technologies. LLMs, trained on vast amounts of text-based human knowledge, offer powerful tools to help scale manual curation efforts. This manuscript describes the results of our initial attempt to adopt LLM-driven AI technologies for this purpose for the Reactome pathway knowledgebase. The approaches and tools described here should be broadly applicable to community knowledgebases.

**Figure 6.**
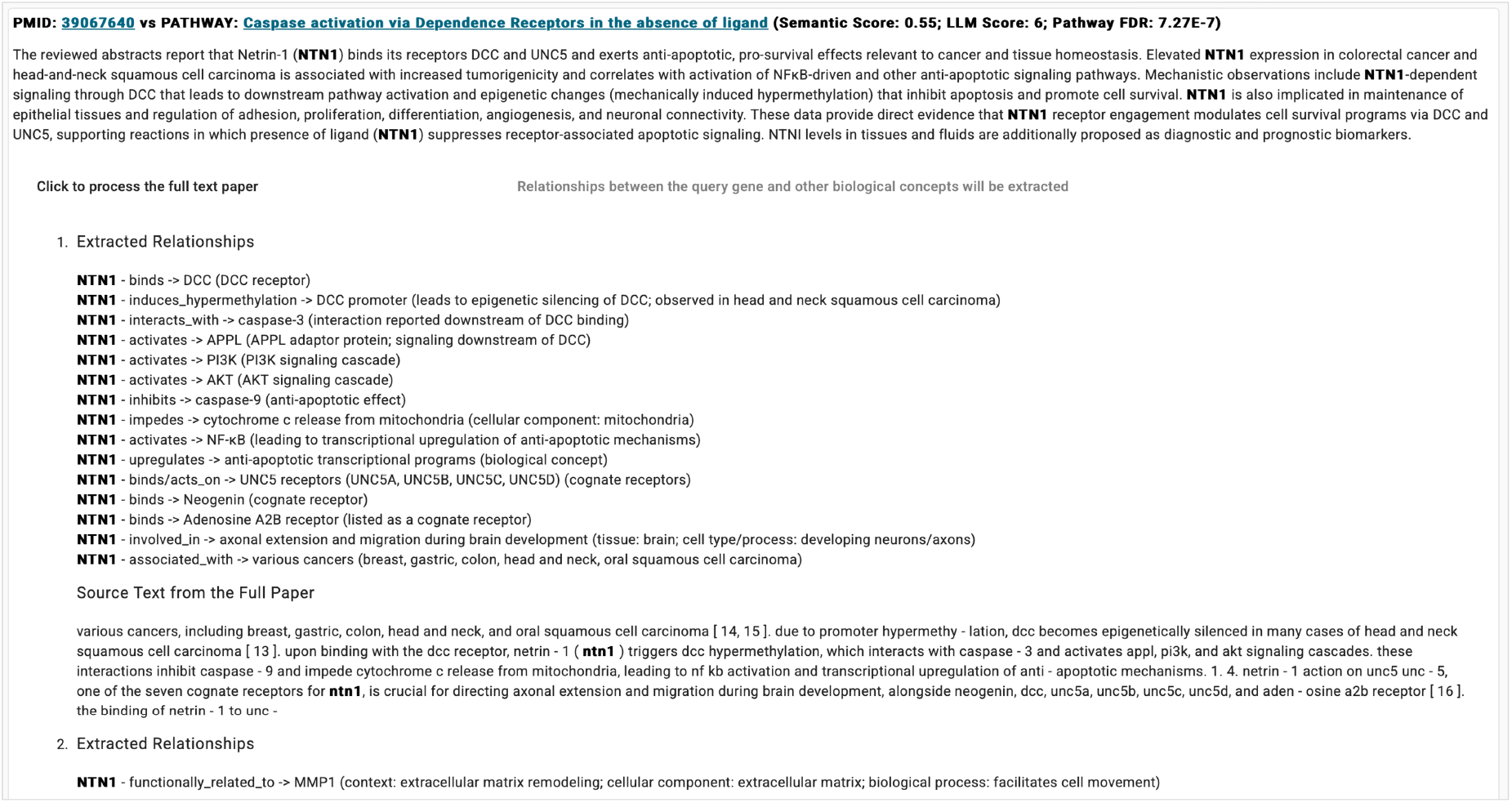
Screenshot of the LLM app in the web-based Reactome curation tool showing functional relationships among biological concepts extracted by the LLM from a full-text PDF.

In this first stage of development, we aimed to create an LLM workflow to assist in annotating human genes to existing Reactome pathways. The workflow is built on a RAG strategy: first predicting interacting pathways for a query gene using protein–protein interactions or functional interactions, then searching PubMed and analyzing the abstracts through semantic similarity scoring and LLM-based judging to obtain relevant literature. Finally, the workflow prompts LLMs to generate text summaries for manual review by Reactome curators. We evaluated the workflow computationally by comparing its predictions with the manual annotations of genes added to Reactome across different releases, demonstrating substantial semantic alignment with curator-derived results. Manual validation by Reactome curators on 19 genes also showed promising outcomes: Over half of the outputs were considered useful as starting points for further refinement. Based on manual validation, we improved the early version of the workflow. To support broader testing and eventual adoption, we also developed an LLM app within the upcoming web-based Reactome curation tool, providing interfaces that allow curators and community users to tune parameters and extract functional relationships from full-text PDF articles.

Manual validation revealed several areas for further improvement. First, the workflow needs to better identify high-quality supporting literature from PubMed by more closely mimicking curators’ strategies for selecting relevant papers (e.g., expanding the keywords used in PubMed searches). Second, the generated text summaries describe overall pathway-level annotations but do not yet conform to Reactome’s object-based data model, which centers on biochemical reactions. In future iterations, we plan to provide structured examples (e.g. “few-shot” prompts) and guide LLMs to generate output aligned with the Reactome data model. Additionally, to better capture biological context (e.g., protein localization, species) and experimental systems (e.g., cell lines, organoids), we will extend the workflow to incorporate full-text papers, maintain additional history for LLMs, and increase text chunk sizes. We will also prompt LLMs to check gene and protein synonyms by providing synonym lists in the context, addressing the synonym issue observed during manual validation for HABP2 (**Table 1**).

The current workflow focuses on annotating genes to existing Reactome pathways. For genes already included in Reactome, the workflow may predict their involvement in additional pathways, thereby revealing pathway crosstalk. For genes not yet annotated, the workflow predicts potential pathways in which they may function, providing biological context that may help reveal functions of understudied or “dark” proteins^18^, similar to efforts in the Reactome IDG portal^7, 8^. We also plan to extend the workflow to support annotation of entirely new pathways by collecting topic (e.g. tumor microenvironment)-related full-text papers and using the LLM workflow to extract functional relationships and automatically generate structured representations consistent with the Reactome data model. Furthermore, the workflow may be expanded to monitor new publications related to already annotated pathways and provide notifications for curators when updates may be needed.

In summary, this manuscript reports the first stage of adopting LLM/AI technologies to support Reactome’s manual curation efforts. The encouraging results give us confidence to further refine and expand the workflow to help scale Reactome curation while maintaining data quality. LLM-driven AI technologies are transforming nearly all scientific fields. Their application in the biomedical research field, including bioinformatics database curation and automatic publication analysis^19^, is still emerging. Recent years have seen rapid progresses, including automated workflows for single-cell RNA-sequencing analysis^20^ and general data-analysis pipelines^21^. The field continues to evolve quickly. As a foundational bioinformatics resource focused on biological pathways, Reactome is actively adopting these new technologies to collect high-quality, well-organized content and provide intuitive tools for the community.

## Methods

### Data sources

Human protein–protein interaction data were downloaded from IntAct in the PSI-MI TAB 2.7 format (https://www.ebi.ac.uk/intact/interactomes, accessed March 2025) and from BioGRID (release 4.4.243, https://downloads.thebiogrid.org/File/BioGRID/Release-Archive/BIOGRID-4.4.243/BIOGRID-ORGANISM-4.4.243.tab3.zip, accessed March 2025). The Reactome functional interaction file (Release 2025), generated by a trained random forest model, was also used. Protein– protein interactions were loaded directly from their original text-based formats, whereas the Reactome functional interactions were first imported into a MongoDB database and then queried via a Python-based API (MongoDB version 8.0.4,https://www.mongodb.com).

To improve the performance of processing PubMed-indexed abstracts, we downloaded the annual PubMed baseline from its FTP site (https://ftp.ncbi.nlm.nih.gov/pubmed/baseline) and loaded all abstracts into a MongoDB database. Abstracts were indexed by PMID to increase search speed. The 2024 baseline, downloaded in March 2025, includes abstracts published through February 2025. In total, we obtained 26,756,819 abstracts indexed by PMID in the MongoDB database.

For development of the LLM workflow, we used a Neo4j graph database converted from a snapshot of the Reactome MySQL curation database (dated June 16, 2025) using the Reactome graph-importer codebase (https://github.com/reactome/graph-importer). To leverage functional interactions for predicting interacting pathways for query genes, we initially used the Reactome IDG portal deployed at https://idg.reactome.org and later ported the code into a Python script to leverage the loaded protein interaction data.

### Implementation of the LLM workflow

We adopted the open-source Python package LangChain (version 0.1.5, https://www.langchain.com) to implement the LLM workflow. To better control performance when querying PubMed-related data, such as interactions, reactions, pathways related to the query gene, and abstracts stored in the local MongoDB database, we implemented a customized retriever, ReactomePubMedRetriever. To compute interacting pathways using protein–protein interactions or Reactome functional interactions, we used the method implemented in the Reactome IDG Portal, which performs pathway enrichment analysis on interacting partners of the query gene using the binomial test. To control for pathway hierarchy in Reactome, only pathways with entity-level views containing reactions were selected as candidates. The Reactome version used for the interacting pathway prediction was Release 91, released in December, 2024.

To calculate the relevance between interacting pathways and abstracts, abstracts were first fetched from the local MongoDB database using PMIDs returned from the PubMed server. Abstracts were then split into text chunks using SentenceTransformersTokenTextSplitter in LangChain, with parameters chunk_overlap=10, model_name=“all-MiniLM-L6-v2”, and add_start_index=True. Each chunk was embedded using SentenceTransformer (sentence-transformers version 2.2.2, https://sbert.net) with the same model. To estimate how likely a publication supports annotation of the query gene to a given interacting pathway, the workflow collected the roles of these interacting partners as annotated in reactions in the interacting pathway, concatenated these descriptions with the pathway summary, and computed the average cosine similarity between this concatenated text and the abstract embeddings. The concatenated text and the abstract were also submitted to the OpenAI API (https://openai.com/api/) to generate an LLM-based relevance score. The top-matched abstracts and pathways, ranked first by cosine similarity and then by LLM score, were sent to the OpenAI API for generation of text summaries.

To extract functional relationships among biological entities from full-text PDF papers, we used PyPDFLoader in langchain_community (version 0.0.17) to load and split PDFs, and then embedded each text chunk into a FAISS vector store (version 1.7.4, https://ai.meta.com/tools/faiss/). From the FAISS vector store, we retrieved the top 12 text chunks semantically related to the query gene’s interactions, reactions, or pathways. These matched chunks were then submitted to the OpenAI API for LLM-based extraction of functional relationships.

The workflow was implemented using Python 3.10.13. The LLM models used via the OpenAI API are gpt-4o-mini for both computational validation and the manual validation and gpt-5-mini for the enhanced version of the LLM app.

### Major prompts used in the LLM workflow

The following prompt templates are used for a variety of tasks. All prompts can be found at: https://github.com/reactome/curator-tool-llm/blob/main/reactome_llm/ReactomePrompts.py.

Prompt template below is used to summarize pathways annotated for a query gene (text enclosed in {} represents placeholders that will be replaced with actual content through parameter passing):

~~~
The gene below has been annotated in multiple pathways described in the context text below in Reactome. Write a summary having about {total_words} words with focus on the molecular functions of the gene in these pathways. Use the context text only for the summary and don’t speculate anything that is not in the text. Make sure to cite the pathway names in the format like this [Pathway_Name] for each sentence. The pathway names are provided in the context. Write a summary sentence at the end to summarize all results.
gene:^14^
context:{annotated_pathways_text}
~~~

Prompt template to summarize the matched abstract and an interacting pathway for the query gene:

~~~
The text in the abstract section is excerpts of scientific papers’ abstracts collected from PubMed and best matched with pathway text below. Write a summary of the abstract text with about {total_words} words to highlight the query_gene and its interaction with interacting_genes, so that Reactome curators can create reactions based on the original papers. The generated text should be based on the abstract text below only, providing evidence showing the possible functions of the query gene in the pathway, {pathway}. Don’t speculate and don’t mention interacting genes if you cannot see them in the abstract text. Don’t just list interacting genes if there is no information.
query_gene:#x007B;query_gene}
interacting_genes:{interacting_genes}
pathway_text:{pathway_text}
abstract:{abstract_text}
~~~

Prompt template to summarize the results for multiple matches between abstracts and interacting pathways for a query gene

~~~
Write a summary based only on the context to summarize the functions of the query gene with about {total_words} words. Highlight the query gene and its interactions with genes in the interacting_genes list. If you cannot see any genes listed in the interacting genes in the context, it is fine not mentioning them. Don’t speculate! Make sure to cite the original context sources, which are provided at the start of each paragraph before “: “, using a format like this [PMID: 123456]. Don’t just list interacting genes if there is no information.
query_gene: {query_gene}
interacting_genes: {interacting_genes}
context: {context}
~~~

Prompt template to extract functional relationships among biological entities from text chunks extract from full text PDF papers

~~~
Extract functional relationships between the query gene specified below and other genes, proteins, or biological concepts from the following document. Output the relationships in the following format: {query_gene} - relationship_type -> other gene or protein or biological concept. If you can find the cellular component, tissue or cell type, or other experimental system related to the extracted relationships, make sure to list them.
query_gene:{query_gene}
document:{document}
~~~

### Development of the LLM app in the web-based Reactome curation tool

We adopted the open source, type script-based Angular framework (version 17.2.4, https://angular.dev) to develop the LLM app in the web-based Reactome curation tool. To support the front-end app, we wrapped the LLM workflow into a RESTful API using flask, an open source Python package (version 3.0.2, https://flask.palletsprojects.com/en/stable/).

### Computational validation

For computational validation, we adopted the LLM workflow to first collect genes that are annotated in Reactome release 90, but not in release 71 and then used release 71 as the basis to run the LLM workflow to generate the LLM predicted text summary as LLM predicted annotations. Curators’ manual annotations were collected from release 90. The generated text was then embedded by SentenceTransformer using the model “all-MiniLM-L6-v2”, as in the LLM workflow. The plot was generated by seaborn (version 0.13.2, https://seaborn.pydata.org).

### Manual validation

Manual validation was conducted by 5 Reactome curators. Genes were selected by individual curators either randomly or because they were on their list of prioritized curation targets. Curators conducted LLM query starting with the default setups, which were used to generate the manual validation results reported in the Results section, and, if no results were obtained, gradually decreasing the stringency until the LLM tool produced a summary. Curators then read the LLM-provided summary and checked the LLM-cited literature references for relevance. Curators recorded their observations (stringency criteria, relevance of LLM-provided summary and literature references), time spent (from the start of search till the analysis of LLM-provided material ended), and occasionally, when the search involved their highly prioritized curation targets, compared the LLM generated results with their own manual curation results, especially comparing the literature references suggested by the LLM with those used to make the actual annotation.

## Code availability

The LLM workflow and the RESTful API wrapping this workflow are hosted at https://github.com/reactome/curator-tool-llm/tree/main and released at Zenodo with doi: 10.5281/zenodo.18001953 (https://zenodo.org/records/18001953). The Angular-based LLM app front-end is hosted as https://github.com/reactome/curator-tool-frontend, as a part of the upcoming Reactome web-based curator tool.

## Acknowledgements

This project was supported by grants from the US National Institutes of Health (U24HG0012198 and U01CA239069). The content described in this publication is solely the responsibility of the authors and does not necessarily represent the official views of the National Institutes of Health.

## References

1. Khatri P, Sirota M, Butte AJ. Ten years of pathway analysis: current approaches and outstanding challenges. PLoS Comput Biol. 2012;8(2):e1002375. Epub 20120223. doi: 10.1371/journal.pcbi.1002375. PubMed PMID: 22383865; PMCID: PMC3285573.

2. Ragueneau E, Gong C, Sinquin P, Sevilla C, Beavers D, Grentner A, Griss J, Hogue GFJ, Li NT, Matthews L, May B, Milacic M, Mohammadi H, Petryszak R, Rothfels K, Shamovsky V, Stephan R, Tiwari K, Weiser J, Wright A, Gillespie M, Wu G, Stein L, Hermjakob H, D’Eustachio P. The Reactome Knowledgebase 2026. Nucleic Acids Res. 2025. Epub 20251118. doi: 10.1093/nar/gkaf1223. PubMed PMID: 41251150.

3. Yu H, Fan L, Li L, Zhou J, Ma Z, Xian L, Hua W, He S, Jin M, Zhang Y, Gandhi A, Ma X. Large Language Models in Biomedical and Health Informatics: A Review with Bibliometric Analysis. J Healthc Inform Res. 2024;8(4):658–711. Epub 20240914. doi: 10.1007/s41666-024-00171-8. PubMed PMID: 39463859; PMCID: PMC11499577.

4. Tiwari K, Matthews L, May B, Shamovsky V, Orlic-Milacic M, Rothfels K, Ragueneau E, Gong C, Stephan R, Li N, Wu G, Stein L, D’Eustachio P, Hermjakob H. ChatGPT usage in the Reactome curation process. bioRxiv. 2023. Epub 20231108. doi: 10.1101/2023.11.08.566195. PubMed PMID: 37986970; PMCID: PMC10659344.

5. Zhang Y, Li Y, Cui L, Cai D, Liu L, Fu T, Huang X, Zhao E, Zhang Y, Chen Y. Siren’s Song in the AI Ocean: A Survey on Hallucination in Large Language Models. arXiv e-prints. 2023: arXiv: 2309.01219.

6. Lewis P, Perez E, Piktus A, Petroni F, Karpukhin V, Goyal N, Küttler H, Lewis M, Yih W-t, Rocktäschel T. Retrieval-augmented generation for knowledge-intensive nlp tasks. Advances in neural information processing systems. 2020;33:9459–74.

7. Beavers D, Brunson T, Sanati N, Matthews L, Haw R, Shorser S, Sevilla C, Viteri G, Conley P, Rothfels K, Hermjakob H, Stein L, D’Eustachio P, Wu G. Illuminate the Functions of Dark Proteins Using the Reactome-IDG Web Portal. Curr Protoc. 2023;3(7):e845. doi: 10.1002/cpz1.845. PubMed PMID: 37467006; PMCID: PMC10399304.

8. Brunson T, Sanati N, Matthews L, Haw R, Beavers D, Shorser S, Sevilla C, Viteri G, Conley P, Rothfels K, Hermjakob H, Stein L, D’Eustachio P, Wu G. Illuminating Dark Proteins using Reactome Pathways. bioRxiv. 2023. Epub 20230605. doi: 10.1101/2023.06.05.543335. PubMed PMID: 37333417; PMCID: PMC10274615.

9. Wu G, Feng X, Stein L. A human functional protein interaction network and its application to cancer data analysis. Genome Biol. 2010;11(5):R53. Epub 20100519. doi: 10.1186/gb-2010-11-5-r53. PubMed PMID: 20482850; PMCID: PMC2898064.

10. Fabregat A, Korninger F, Viteri G, Sidiropoulos K, Marin-Garcia P, Ping P, Wu G, Stein L, D’Eustachio P, Hermjakob H. Reactome graph database: Efficient access to complex pathway data. PLoS Comput Biol. 2018;14(1):e1005968. Epub 20180129. doi: 10.1371/journal.pcbi.1005968. PubMed PMID: 29377902; PMCID: PMC5805351.

11. UniProt C. UniProt: the Universal Protein Knowledgebase in 2025. Nucleic Acids Res. 2025;53(D1):D609–D17. doi: 10.1093/nar/gkae1010. PubMed PMID: 39552041; PMCID: PMC11701636.

12. Orchard S, Ammari M, Aranda B, Breuza L, Briganti L, Broackes-Carter F, Campbell NH, Chavali G, Chen C, del-Toro N, Duesbury M, Dumousseau M, Galeota E, Hinz U, Iannuccelli M, Jagannathan S, Jimenez R, Khadake J, Lagreid A, Licata L, Lovering RC, Meldal B, Melidoni AN, Milagros M, Peluso D, Perfetto L, Porras P, Raghunath A, Ricard-Blum S, Roechert B, Stutz A, Tognolli M, van Roey K, Cesareni G, Hermjakob H. The MIntAct project--IntAct as a common curation platform for 11 molecular interaction databases. Nucleic Acids Res. 2014;42(Database issue):D358–63. Epub 20131113. doi: 10.1093/nar/gkt1115. PubMed PMID: 24234451; PMCID: PMC3965093.

13. Ashburner M, Ball CA, Blake JA, Botstein D, Butler H, Cherry JM, Davis AP, Dolinski K, Dwight SS, Eppig JT, Harris MA, Hill DP, Issel-Tarver L, Kasarskis A, Lewis S, Matese JC, Richardson JE, Ringwald M, Rubin GM, Sherlock G. Gene ontology: tool for the unification of biology. The Gene Ontology Consortium. Nat Genet. 2000;25(1):25–9. doi: 10.1038/75556. PubMed PMID: 10802651; PMCID: PMC3037419.

14. Gene Ontology C, Aleksander SA, Balhoff J, Carbon S, Cherry JM, Drabkin HJ, Ebert D, Feuermann M, Gaudet P, Harris NL, Hill DP, Lee R, Mi H, Moxon S, Mungall CJ, Muruganugan A, Mushayahama T, Sternberg PW, Thomas PD, Van Auken K, Ramsey J, Siegele DA, Chisholm RL, Fey P, Aspromonte MC, Nugnes MV, Quaglia F, Tosatto S, Giglio M, Nadendla S, Antonazzo G, Attrill H, Dos Santos G, Marygold S, Strelets V, Tabone CJ, Thurmond J, Zhou P, Ahmed SH, Asanitthong P, Luna Buitrago D, Erdol MN, Gage MC, Ali Kadhum M, Li KYC, Long M, Michalak A, Pesala A, Pritazahra A, Saverimuttu SCC, Su R, Thurlow KE, Lovering RC, Logie C, Oliferenko S, Blake J, Christie K, Corbani L, Dolan ME, Drabkin HJ, Hill DP, Ni L, Sitnikov D, Smith C, Cuzick A, Seager J, Cooper L, Elser J, Jaiswal P, Gupta P, Jaiswal P, Naithani S, Lera-Ramirez M, Rutherford K, Wood V, De Pons JL, Dwinell MR, Hayman GT, Kaldunski ML, Kwitek AE, Laulederkind SJF, Tutaj MA, Vedi M, Wang SJ, D’Eustachio P, Aimo L, Axelsen K, Bridge A, Hyka-Nouspikel N, Morgat A, Aleksander SA, Cherry JM, Engel SR, Karra K, Miyasato SR, Nash RS, Skrzypek MS, Weng S, Wong ED, Bakker E, Berardini TZ, Reiser L, Auchincloss A, Axelsen K, Argoud-Puy G, Blatter MC, Boutet E, Breuza L, Bridge A, Casals-Casas C, Coudert E, Estreicher A, Livia Famiglietti M, Feuermann M, Gos A, Gruaz-Gumowski N, Hulo C, Hyka-Nouspikel N, Jungo F, Le Mercier P, Lieberherr D, Masson P, Morgat A, Pedruzzi I, Pourcel L, Poux S, Rivoire C, Sundaram S, Bateman A, Bowler-Barnett E, Bye AJH, Denny P, Ignatchenko A, Ishtiaq R, Lock A, Lussi Y, Magrane M, Martin MJ, Orchard S, Raposo P, Speretta E, Tyagi N, Warner K, Zaru R, Diehl AD, Lee R, Chan J, Diamantakis S, Raciti D, Zarowiecki M, Fisher M, James-Zorn C, Ponferrada V, Zorn A, Ramachandran S, Ruzicka L, Westerfield M. The Gene Ontology knowledgebase in 2023. Genetics. 2023;224(1). doi: 10.1093/genetics/iyad031. PubMed PMID: 36866529; PMCID: PMC10158837.

15. Oughtred R, Rust J, Chang C, Breitkreutz BJ, Stark C, Willems A, Boucher L, Leung G, Kolas N, Zhang F, Dolma S, Coulombe-Huntington J, Chatr-Aryamontri A, Dolinski K, Tyers M. The BioGRID database: A comprehensive biomedical resource of curated protein, genetic, and chemical interactions. Protein Sci. 2021;30(1):187–200. Epub 20201123. doi: 10.1002/pro.3978. PubMed PMID: 33070389; PMCID: PMC7737760.

16. Zheng L, Chiang W-L, Sheng Y, Zhuang S, Wu Z, Zhuang Y, Lin Z, Li Z, Li D, Xing E. Judging llm-as-a-judge with mt-bench and chatbot arena. Advances in neural information processing systems. 2023;36:46595–623.

17. Malik A, Arsalan M, Moreno C, Mosquera J, Felix E, Kiziloren T, Muthukrishnan V, Zdrazil B, Leach AR, O’Boyle NM. ChEBI: re-engineered for a sustainable future. Nucleic Acids Res. 2025. Epub 20251128. doi: 10.1093/nar/gkaf1271. PubMed PMID: 41312627.

18. Oprea TI. Exploring the dark genome: implications for precision medicine. Mamm Genome. 2019;30(7-8):192–200. Epub 20190704. doi: 10.1007/s00335-019-09809-0. PubMed PMID: 31270560; PMCID: PMC6836689.

19. Lála J, O’Donoghue O, Shtedritski A, Cox S, Rodriques SG, White AD. Paperqa: Retrieval-augmented generative agent for scientific research. arXiv preprint arXiv:231207559. 2023.

20. Alber S, Chen B, Sun E, Isakova A, Wilk AJ, Zou J. CellVoyager: AI CompBio Agent Generates New Insights by Autonomously Analyzing Biological Data. bioRxiv. 2025:2025.06.03.657517. doi: 10.1101/2025.06.03.657517.

21. Huang K, Zhang S, Wang H, Qu Y, Lu Y, Roohani Y, Li R, Qiu L, Li G, Zhang J, Yin D, Marwaha S, Carter JN, Zhou X, Wheeler M, Bernstein JA, Wang M, He P, Zhou J, Snyder M, Cong L, Regev A, Leskovec J. Biomni: A General-Purpose Biomedical AI Agent. bioRxiv. 2025. Epub 20250602. doi: 10.1101/2025.05.30.656746. PubMed PMID: 40501924; PMCID: PMC12157518.

